# rpSTS Stimulation Disrupts Shared Neural Responses to Social Content during Naturalistic Viewing

**DOI:** 10.64898/2026.10.08.757709

**Authors:** James C Thompson, Courtney Marsh, Peter Kakalec, Katie Rawlings, Rebecca Roy

## Abstract

Shared understanding of common experiences can foster feelings of social closeness and friendship, and help us understand those experiences. Recent work has suggested that patterns of neural activity shared across individuals, especially throughout the default mode network, may underlie shared representations. Here we used repetitive transcranial magnetic stimulation (rTMS) and functional magnetic resonance imaging during movie watching to show that shared neural patterns to common experiences rely on the encoding of social information from the environment by the right posterior superior temporal sulcus (rpSTS). An encoding model of movie features showed that rTMS selectively degraded social feature encoding in rpSTS, leaving low-level visual encoding in early visual cortex intact. Using natural language processing (NLP)-based analysis, we found that rpSTS stimulation was associated with reduced memory and intersubject pattern similarity across DMN and limbic networks during scenes central to the movie narrative. Projecting scene embeddings onto NLP-based concept axes revealed reductions in memory and neural pattern similarity for scenes conveying helplessness and empathy following rTMS. These findings provide causal evidence that the rpSTS is a key region that encodes shared social information into common neural representations, and suggest that disrupting its input leads DMN representations to become idiosyncratic.

## Introduction

Seeing the world in a similar way to others is critical for how we form, maintain, and deepen our social relationships. Humans are highly motivated to experience and share commonalities in our inner states with others [1,2]. Shared representations of the world can help make us closer to others. They also provide a means for us to verify with others what is real [1,3]. Shared neural activity when experiencing the same events has been linked to a shared understanding between individuals [4,5], and is associated with the likelihood that two individuals will become friends [6,7]. Shared patterns of neural activity in the default mode network (DMN) during naturalistic viewing have been argued to be especially important to the formation of shared representations [8,9,10]. However, it is not clear how the common experience of the external world is transformed into a shared neural representation in higher order networks such as the DMN. One theory is that the DMN integrates perceptual information with idiosyncratic internal representations built on long-term memories and perspectives [8,10]. Given this idea, we hypothesized that degrading the neural encoding of information from the environment might decrease shared representations in DMN and related networks, leading to more idiosyncratic neural activity.

While shared representations are unlikely to be exclusively social in content, they might be particularly important for understanding social information, which is often ambiguous and not directly observable. Social information is a key driver of shared neural responses in the DMN [11,12]. The right posterior superior temporal sulcus (rpSTS) is one region that might provide socially relevant external information to the DMN and other higher-order cortical regions. The rpSTS preferentially responds to dynamic, multimodal social information [13], and has strong anatomical and functional connections to and from dorsal and ventral visual pathways, as well as DMN and limbic regions including the amygdala [14–22]. Previous work shows lesions to superior temporal cortex lead to deficits in social perception [23], and reversible disruption of rpSTS activity via repetitive transcranial magnetic stimulation (rTMS) produces impairments in the recognition of social and emotional stimuli [24,25]. While the rpSTS has been argued to be an important hub region within larger networks (including the DMN) that process social information [11], there is no direct evidence that manipulating activity in this region, such as via neuromodulation, leads to changes in shared neural representations in the DMN.

In the present study, we used continuous theta-burst rTMS and fMRI to examine the role of the rpSTS in the spontaneous encoding of social information and shared representations during naturalistic viewing. We predicted that the disruption of neural activity in rpSTS by rTMS would impair the encoding of social information during movie watching, leading to decreased shared representations and more idiosyncratic processing in networks such as the DMN.

## Results

Participants (n=41) first attended a scanning session in which we acquired BOLD fMRI and anatomical data as participants viewed a functional localizer task consisting of blocks of videos of moving people, scenes, objects, food, and scrambled versions of these videos (Figure 1A). TMS resting motor thresholds (RMTs) were also acquired for each participant. In a subsequent session, participants returned and received continuous theta-burst (cTBS) TMS to either the functionally localized rpSTS (rpSTS group; n=21, 17F) or a vertex control location (vertex group; n=20, 14F). Immediately following cTBS TMS administration participants entered the scanner and we acquired BOLD fMRI data as participants viewed a short (20min) movie (“The Neighbors’ Window”). Upon completion of scanning, participants freely recalled details from the movie and completed a recognition memory test with test items related to the movie content.

**Figure 1.**
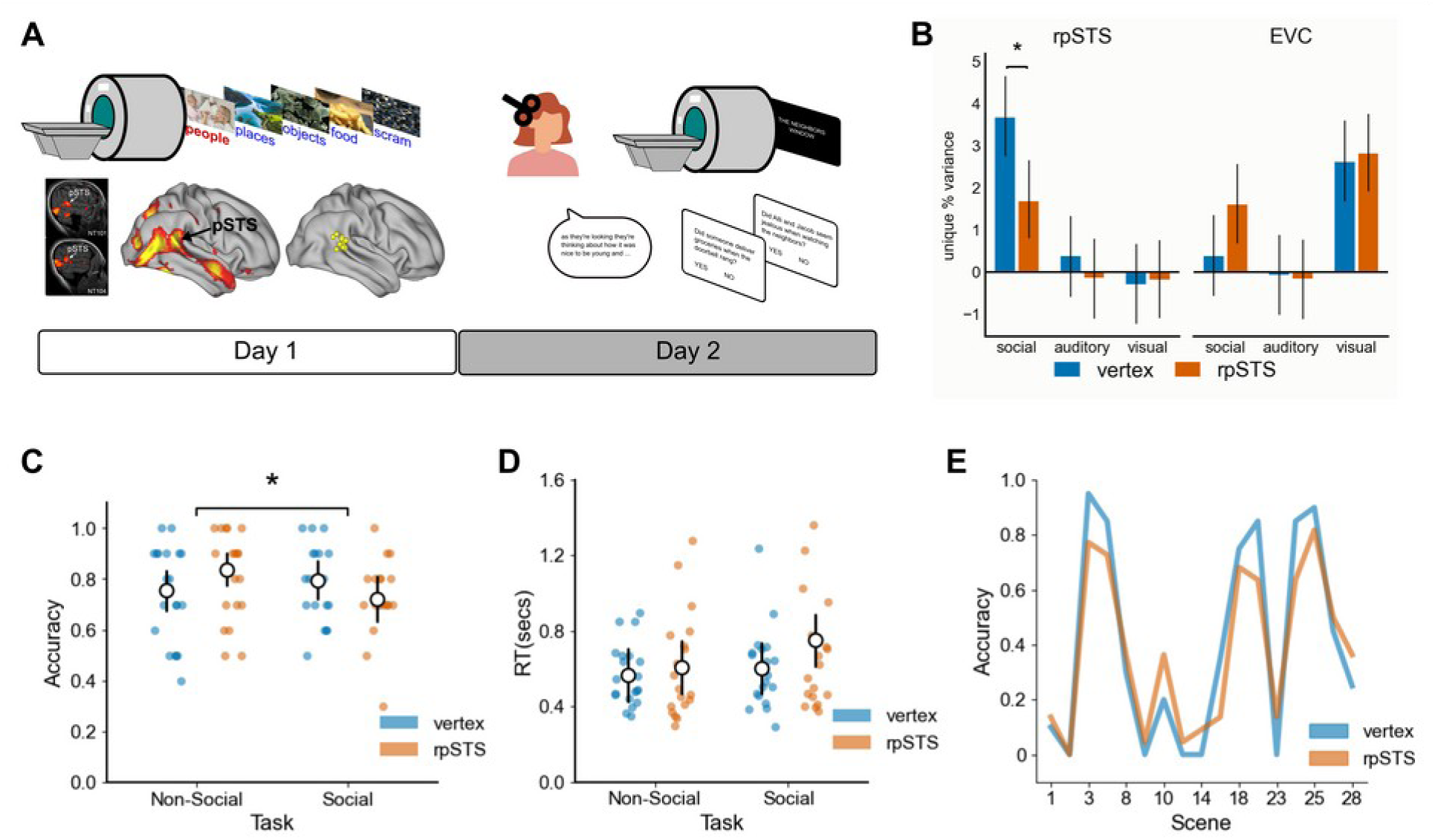
Schematic of study timeline and effects of rTMS on neural encoding of movie features and memory for movie content. **A.** On day 1, participants underwent MRI scanning as they viewed a functional localizer task to identify rpSTS. Individualized rpSTS loci were used as targets for rTMS. On day 2, participants received continuous theta-burst rTMS to rpSTS or vertex and then underwent MRI scanning as they viewed a movie. Following the movie, participants freely recalled as many details from the movie as they could and performed a recognition memory test. **B.** The encoding model prediction performance (posterior mean +/- 95% HDI unique explained variance for each group of model features) from the rpSTS and EVC regions of interest. Unique variance of the social features was reduced by rTMS to the rpSTS (red) relative to rTMS to the vertex (blue) in the rpSTS ROI (left), while unique variance of the visual features was not reduced in the EVC (right). Line and asterisk indicate group posterior means with non-overlapping 95% HDIs. **C.** Participants showed an interaction between stimulation group (rpSTS: red; or vertex: blue) and content (social or non-social) in recognition memory accuracy. Line and asterisk indicate interaction posterior where 95% HDI does not contain zero. **D.** Participants did not show differences in recognition memory RT as a function of stimulation group or content. **E.** Scene recall from each scene, showing that there was greater recall for scenes at the start and end of the movie, relative to scenes in the middle, for both groups.

### rTMS Disrupts Encoding of Social Features in rpSTS

We first sought to verify that rTMS disrupted the neural encoding of social information during movie viewing. One challenge to analyzing naturalistic stimuli is that there are not clear boundaries between conditions of interest, such as social versus nonsocial scenes in the movie. To address this challenge, we fit an encoding model to the fMRI response during movie watching from an rpSTS region of interest (ROI), and examined if the encoding of social features was selectively reduced following rTMS to either the rpSTS or a vertex control site. We used computer vision methods to quantify social features, such as the number of faces, number of bodies, and the amount of biological motion, throughout the movie. We also extracted the amount of speech, and had independent judges rate the degree of communicative interactions, on a moment-by-moment basis. To rule out a nonselective perceptual disruption, we also quantified visual features (motion energy) and auditory features (volume amplitude fluctuations) throughout the movie, and to exclude a more general effect (e.g., arousal) of rTMS we compared the rpSTS to the response from an early visual cortex (EVC) ROI.

Using cross-validated ridge regression and testing the prediction performance of the encoding model on held-out segments of data, we found that stimulation of the rpSTS reduced overall model prediction performance relative to vertex stimulation in the rpSTS ROI (rpSTS group: r = 0.08 vs vertex group: r = 0.17) but not the EVC ROI (rpSTS group: r = 0.23 vs vertex group: r = 0.20), posterior mean *b* = -0.13, 95% HDI = [-0.21, -0.05]. Importantly, we also observed an interaction between stimulation group, ROI, and features when we examined unique explained variance of each group of features, such that rpSTS stimulation was associated with lower encoding of social features relative to the visual or auditory features in the rpSTS ROI, but not the EVC ROI, when compared to the vertex group, posterior mean *b* = -0.03, 95% HDI = [-0.05, - 0.01] (Figure 1B). In contrast, there were no differences in the encoding of low-level visual features in the EVC ROI between the rpSTS and vertex groups, and auditory features did not consistently predict unique variance in either rpSTS or EVC ROI. These results indicate that rTMS decreased the processing of social information in the rpSTS, and that this decrease was not due to either a general perceptual decrease or nonspecific effects.

### rTMS to the rpSTS and Recognition Memory for Social Content

We then examined if rpSTS disruption impaired memory for social details from the movie. After the movie was completed, participants performed a recognition memory test that included items related to the social content and items related to nonsocial content of the movie (Figure 1A). Looking first at recognition accuracy for the vertex and rpSTS groups, we observed an interaction between group and movie content, posterior mean *b =* -0.93, 95% HDI=[-1.74, -0.11]. This interaction was driven by lower recognition accuracy for social content relative to nonsocial content for the rpSTS group, posterior mean *difference* = -0.09, 95% HDI=[-0.18, -0.01], but not the vertex group posterior mean *difference* = 0.04, 95% HDI=[-0.04, 0.13] (Figure 1C). This finding is consistent with a decrease in the encoding of social information following disruption of rpSTS activity, perhaps resulting in better encoding of nonsocial details. No interaction between stimulation groups and movie content was observed for recognition memory (log transformed) reaction times, posterior mean *b =* 0.11, 95% HDI=[-0.05, 0.28] (Figure 1D).

To analyze free recall, we first divided the movie into distinct scenes based on ratings from independent raters. Some scenes were brief (<3secs) transitions and were excluded from recall analysis. For the remaining 17 scenes, we identified from free recall data if a participant recalled a scene. The proportion of participants in each group who recalled a scene, for all scenes, is shown in Figure 1E. To assess whether recall varied across scenes, between vertex and rpSTS groups, or their interaction, we compared nested Bayesian regression models using approximate leave-one-out cross-validation (LOO-CV). Including scene substantially improved predictive fit relative to a model that tested recall regardless of scene (ΔELPD = 201.3, SE = 15.9), indicating that recall differed across scenes. Adding group to the scene model did not improve predictive fit (ΔELPD = 0.1, SE = 0.8). Including the group × scene interaction led to a small increase in model fit (ΔELPD = 6.1, SE = 4.2), but not enough to strongly support the conclusion that there was an interaction between scene and stimulation group. The recall curves from the two groups resembled serial position effects, and indeed across both groups the scene recall from the first third of the movie was greater than scene recall from the middle third, posterior mean *b =* -1.15, 95% HDI=[-1.77, -0.57], but not the final third, posterior mean *b =* 0.32, 95% HDI=[-0.23, 0.92]. However, for the recall of the naturalistic movies such as the one used in this study it can be difficult to disentangle memory-based serial position effects from effects driven by the movie narrative. The following analyses sought to clarify the contribution of the narrative to recall and shared neural responses.

### rTMS to the rpSTS Alters Intersubject Synchrony and Pattern Similarity During Naturalistic Viewing

We examined the effects of rTMS to rpSTS on shared neural representations first by calculating intersubject correlations (ISC) of fMRI time series from 400 cortical parcels and 16 subcortical parcels. Decreased ISC in the rpSTS group relative to the vertex group was observed in right inferior frontal, orbitofrontal, somatomotor, and anterior temporal cortices, left anterior intraparietal, midcingulate, and superior temporal cortices, and in the left amygdala (p<0.05 FDR corr; Figure 2A,B). In contrast, we also saw increased ISC in the rpSTS group relative to the vertex group in multiple extrastriate cortical regions and bilateral posterior insula (p<0.05 FDR corr). To examine effects of rpSTS on ISC at the network-level, we averaged ISC estimates within regions of a 17-node network parcellation. These data showed that the rpSTS group had higher ISC in Visual-Central and Visual-Peripheral nodes relative to the vertex group (p<0.05 FDR corr; Figure 2C). Together, these results suggest that rTMS to rpSTS decreased neural synchrony in higher order regions downstream to rpSTS, while neural synchrony in lower order visual regions upstream to rpSTS was increased, relative to vertex.

**Figure 2.**
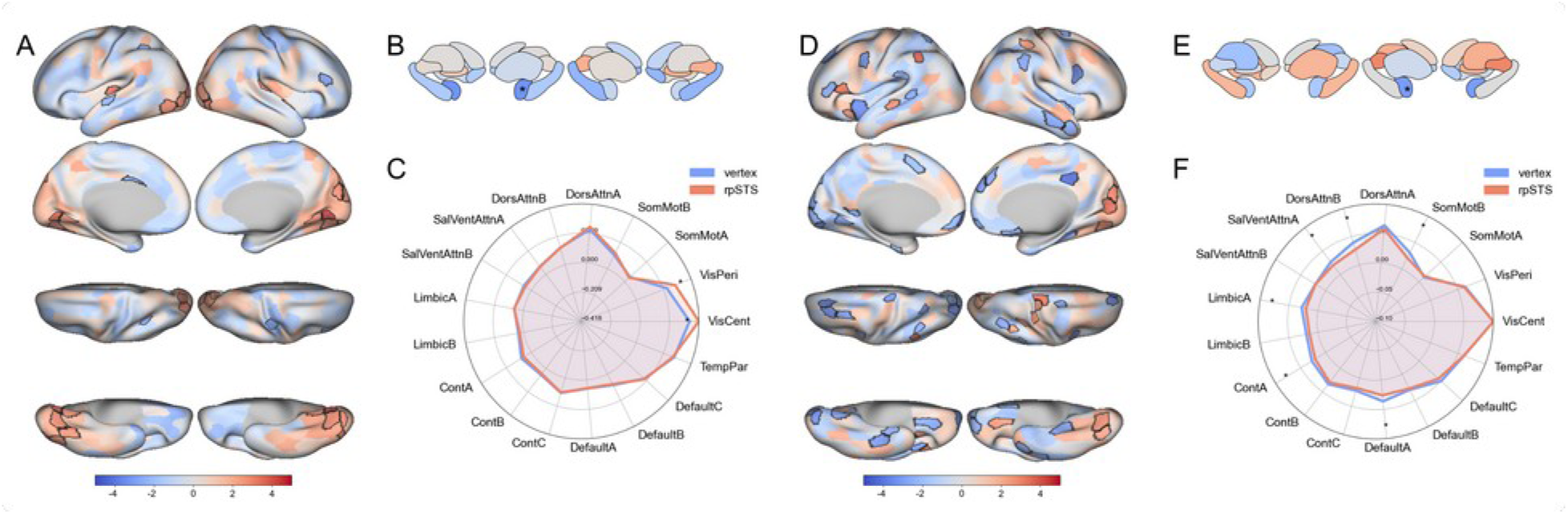
The effects of rTMS to rpSTS versus vertex on neural synchrony and pattern similarity. **A & B.** Differences between the rpSTS and vertex groups in intersubject synchrony (ISC) during movie watching. The t-values of the comparison of intragroup ISC are shown, with lower ISC in the rpSTS relative to vertex shown in cool colors, and higher ISC in the rpSTS relative to vertex group shown in warm colors. Parcels are overlaid on the fsaverage6 cortical surface **(A)** or the Melbourne Subcortex Atlas (**B**). Parcels outlined in black or overlaid with an asterisk are p<0.05 (FDR corr). C. Data from the 400 cortical parcels collapsed into 17 networks, showing mean within group ISC for the rpSTS group (red) and the vertex group (blue). Asterisks indicate significant differences between the groups, p<0.05 (FDR corr). **D & E.** Differences between the rpSTS and vertex groups in intersubject pattern similarity (ISPS) during movie watching averaged across all scenes. The t-values of the comparison of intragroup ISPS are shown, with lower ISPS in the rpSTS relative to vertex shown in cool colors, and higher ISPS in the rpSTS relative to vertex group shown in warm colors. F. Data from the 400 cortical parcels collapsed into 17 networks, showing mean within group ISPS for the rpSTS group (red) and the vertex group (blue). Asterisks indicate significant differences between the groups, p<0.05 (FDR corr).

To further examine the effects of rpSTS versus vertex rTMS on shared neural representations, we examined multivoxel patterns of activity from the 416 cortical and subcortical parcels for each participant within each movie scene, and calculated the intersubject pattern similarity (ISPS) between participants within each group for all the scenes. These data revealed that ISPS was lower in the rpSTS group, compared to the vertex group, in right anterior temporal, temporoparietal, inferior frontal, dorsomedial prefrontal, orbitofrontal, and midcingulate cortices, and right amygdala, as well as left midSTS, anterior temporal, extrastriate visual, premotor, inferior frontal, and anterior insula cortices (p<0.05 FDR corr; Figure 2D,E). Conversely, we saw higher ISPS in the rpSTS group in right extrastriate and superior parietal cortices, as well as left midSTS, temporoparietal, inferior frontal, and anterior insula cortices, relative to the vertex group (p<0.05 FDR corr). At the network level, reduced ISPS was observed in Somatomotor-B, Dorsal Attention-A, Control-A, Limbic-A, and Default-A Networks in the rpSTS group compared to the vertex group (Figure 2F).

### Recall of Central Scenes is Lower Following rpSTS Stimulation

While there was no overall difference between rpSTS and vertex groups in the probability of recalling a scene, results from the recognition memory task and the ISC and ISPS data suggest that the rpSTS group might have encoded the movie stimulus differently from the vertex group. We therefore examined if different content within the movie was used by each group during movie encoding. Emotional arousal, in particular, has been shown to be an important contributor to the subsequent recall of scene details from naturalistic stimuli [26,27]. Additionally, scene-level ratings consider each scene in isolation but do not capture how scenes relate to each other or the overall narrative. Recent network-based approaches to understanding narrative memory have shown that scenes central to the narrative are recalled at a higher rate than noncentral scenes [28]. We used scene-level annotations of the movie to obtain semantic embeddings for each scene using a sentence-level contextual language model (SBERT). We then calculated the cosine similarity of scene embeddings and derived the degree centrality, or how similar a scene is to every other scene, for each scene in the movie (Figure 3A).

**Figure 3.**
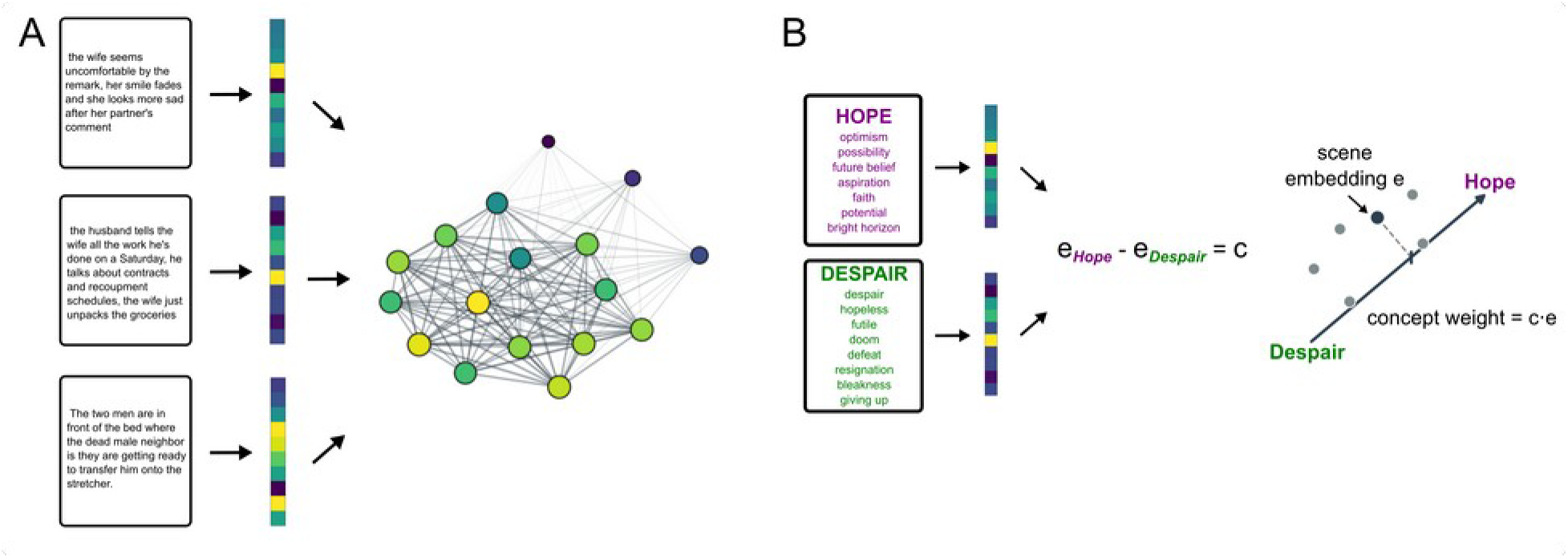
Diagram outlining the scene centrality and conceptual embedding analyses. **A.** Scene transcripts were projected onto 768-dimensional embeddings using SBERT. Embeddings from each scene were normalized and the cosine similarity between all scene embeddings calculated. Degree centrality (i.e., the sum similarity of a scene to all other scenes normalized by dividing by n scenes − 1) was then calculated. **B.** LLM-generated concept descriptions were projected onto the same 768-dimensional embeddings using SBERT. The embeddings from one concept pole were then subtracted from the embedding from the other pole to create a concept axis. Scene embeddings were projected onto these concept axes by finding the cosine similarity to provide a concept weight for that scene.

We observed that probability of scene recall was higher for longer scenes than shorter scenes for both stimulation groups, posterior mean *b* = 0.99, 95% HDI=[0.57, 1.39]. After controlling for scene length, we observed that for both groups higher emotional arousal of a scene was associated with greater probability of scene recall, posterior mean *b =* 1.37, 95% HDI=[0.88, 1.89], and that scene degree centrality was associated with higher probability of scene recall, posterior mean *b =* 1.09, 95% HDI=[0.28, 2.04] (Figure 4A). However, we also observed an interaction between group and degree centrality on the probability of scene recall, posterior mean *b =* -0.90, 95% HDI=[-1.93, - 0.04]. Participants who received rpSTS stimulation were considerably less likely to rely on the centrality of a scene for recall than those in the vertex group. While the recall of the vertex group was biased towards scenes more central to the movie narrative, recall in the rpSTS group appeared to reflect a more even encoding of both central and noncentral scenes (Figure 4A). These results are consistent with the proposal that rTMS disruption of the rpSTS was associated with reduced encoding of the core narrative features of the movie.

**Figure 4.**
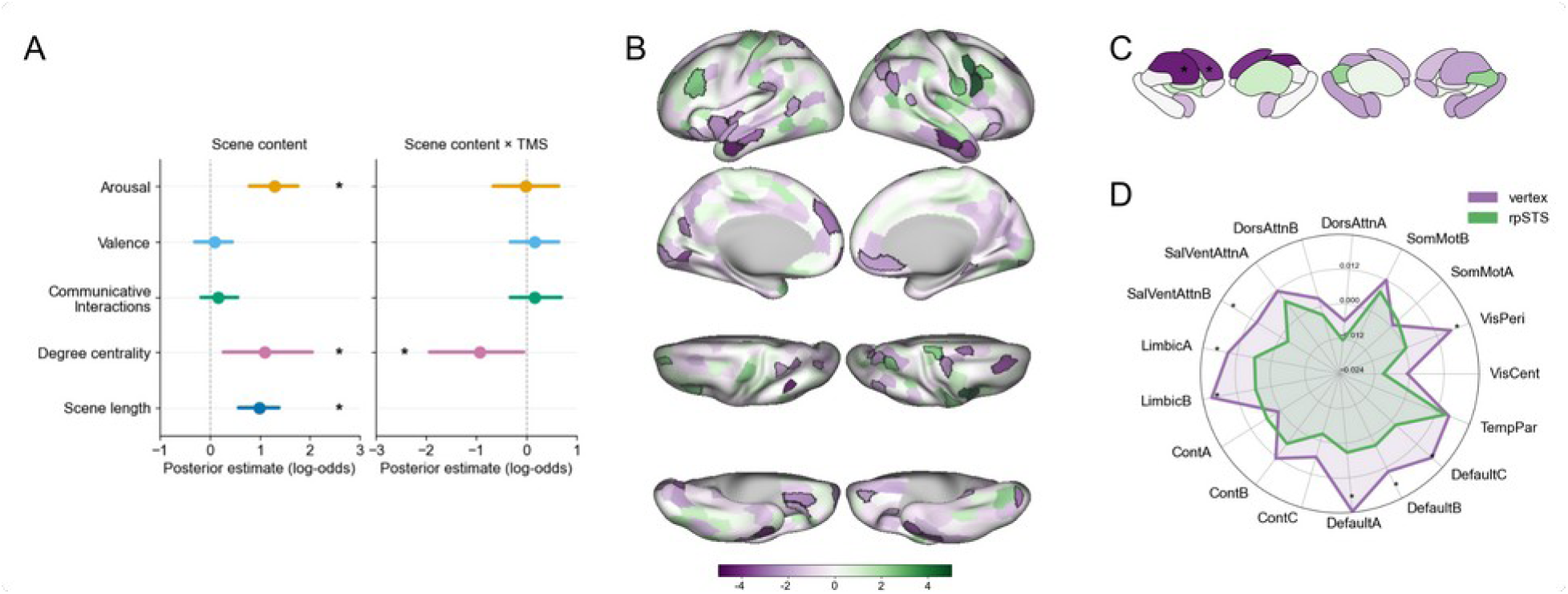
Reduced encoding of central scenes following rTMS to the rpSTS. **A.** Plots represent the posterior mean and 95% HDI estimates of the effects of scene content and rTMS on the probability of scene recall. The main effect of scene content is shown on the left, with the interaction between content and stimulation group on the right. Asterisks indicate that the 95% HDI did not contain zero. **B & C.** Group differences in the difference in ISPS between high and low centrality scenes. The t-values of the comparison between the rpSTS and vertex groups in ISPS from high – low centrality scenes are shown, with lower ISPS difference in the rpSTS relative to vertex shown in purples, and higher ISPS difference in the rpSTS relative to vertex group shown in green. Parcels are overlaid on the fsaverage6 cortical surface **(B)** or the Melbourne Subcortex Atlas (**C**). Parcels outlined in black or overlaid with an asterisk are p<0.05 (FDR corr). **D.** Data from the 400 cortical parcels collapsed into 17 networks, showing mean difference in ISPS between high and low centrality scenes for the rpSTS group (green) and the vertex group (purple). Asterisks indicate significant differences between the groups, p<0.05 (FDR corr).

### Shared Neural Representations of Central Scenes Are Decreased Following rpSTS Stimulation

To examine if rpSTS and vertex groups differed in the neural encoding of scenes according to scene centrality, we examined differences in ISPS for low and high degree centrality scenes between the two groups. As is shown in Figure 4B,C, the rpSTS group showed a smaller difference in ISPS between high and low centrality scenes, compared to the vertex group, in a range of social/affective and default mode regions, including bilateral pSTS, right temporoparietal junction, anterior temporal, orbitofrontal and ventromedial prefrontal cortices, left rostral prefrontal, dorsomedial prefrontal, anterior insula, superior parietal, and medial occipital cortices, and left caudate and putamen (p<0.05 FDR corr). Greater ISPS differences to high versus low centrality scenes in the rpSTS group was observed in right somatomotor and premotor cortices and left dorsolateral prefrontal cortex, relative to the vertex group (p<0.05 FDR corr). At the network level, we observed lower ISPS differences to high versus low centrality scenes in Limbic-A and Limbic-B, Default-A, Default-B, and Default-C, and Salience/VentralAttention-B and Visual-Peripheral sub-networks in the rpSTS group relative to vertex (p<0.05 FDR corr)(Figure 4D). These findings indicate that rpSTS stimulation is associated with reduced shared representations of central narrative scenes in limbic networks and the DMN.

### Narrative Concept Embedding and Recall

While our results indicate that disrupting rpSTS activity via rTMS may reduce the contribution of narrative centrality to scene recall, and is associated with reduced shared neural representations in social/affective and default mode network regions associated with central scenes, they do not provide details of the semantic contents of the narrative key to shared representations. To understand the narrative contents in more detail, we developed a novel method borrowing from NLP approaches to narrative comprehension [29–31] (Figure 3B). We started by identifying several core concepts from film and narrative theory relevant to the movie, with positive and negative poles: danger/safety, chaos/order, agency/helplessness, moral clarity/moral ambiguity, transformation/stasis, joy/grief, hope/despair. We then used a generative LLM (Claude Sonnet 4.6) to generate phrases associated with the positive and negative poles of these concepts, and obtained embeddings for the phrases using SBERT. Subtracting the mean of the negative pole embedding from the mean of the positive pole embedding for each concept yielded a concept embedding. This allowed us to project each scene embedding onto the concept embedding by calculating the cosine similarity, providing a concept weight for which a negative weight reflects similarity to the negative pole of the concept and a positive weight reflects similarity to the positive pole direction.

To examine the contribution of these concept weights to the central narrative of the movie, we first correlated each scene’s concept weights with scene degree centrality. Only moral clarity/moral ambiguity, posterior mean *b =* -0.5, 95% HDI = [-1.01, -0.1], agency/helplessness, posterior mean *b =* -0.44, 95% HDI = [-0.99, -0.03], and hope/despair, posterior mean *b =* -0.76, 95% HDI = [-1.15, -0.41], showed evidence of correlation with scene degree centrality. These results indicated that the central scenes of the movie reflect the negative pole of these concepts (e.g., moral ambiguity, helplessness, despair). We then examined if the concept weights that showed correlations with scene degree centrality were associated with scene recall, and if the rpSTS and vertex groups differed in the extent to which the concept weight was associated with recall. Of the three concept weights, only the agency/helplessness concept was associated with scene recall, posterior mean *b =* -0.66, 95% HDI = [-1.00, - 0.34]. Overall, the higher the helplessness weight of a scene (i.e., the negative pole of the agency/helplessness concept), the better participants recalled the scene (Figure 5A). Furthermore, there was a difference between groups in the extent to which the helplessness content of a scene contributed to recall, posterior mean *b =* 0.42, 95% HDI = [0.02, 0.85]. That is, participants in the rpSTS group used the helplessness concept in a scene less during movie encoding than did the vertex group.

**Figure 5.**
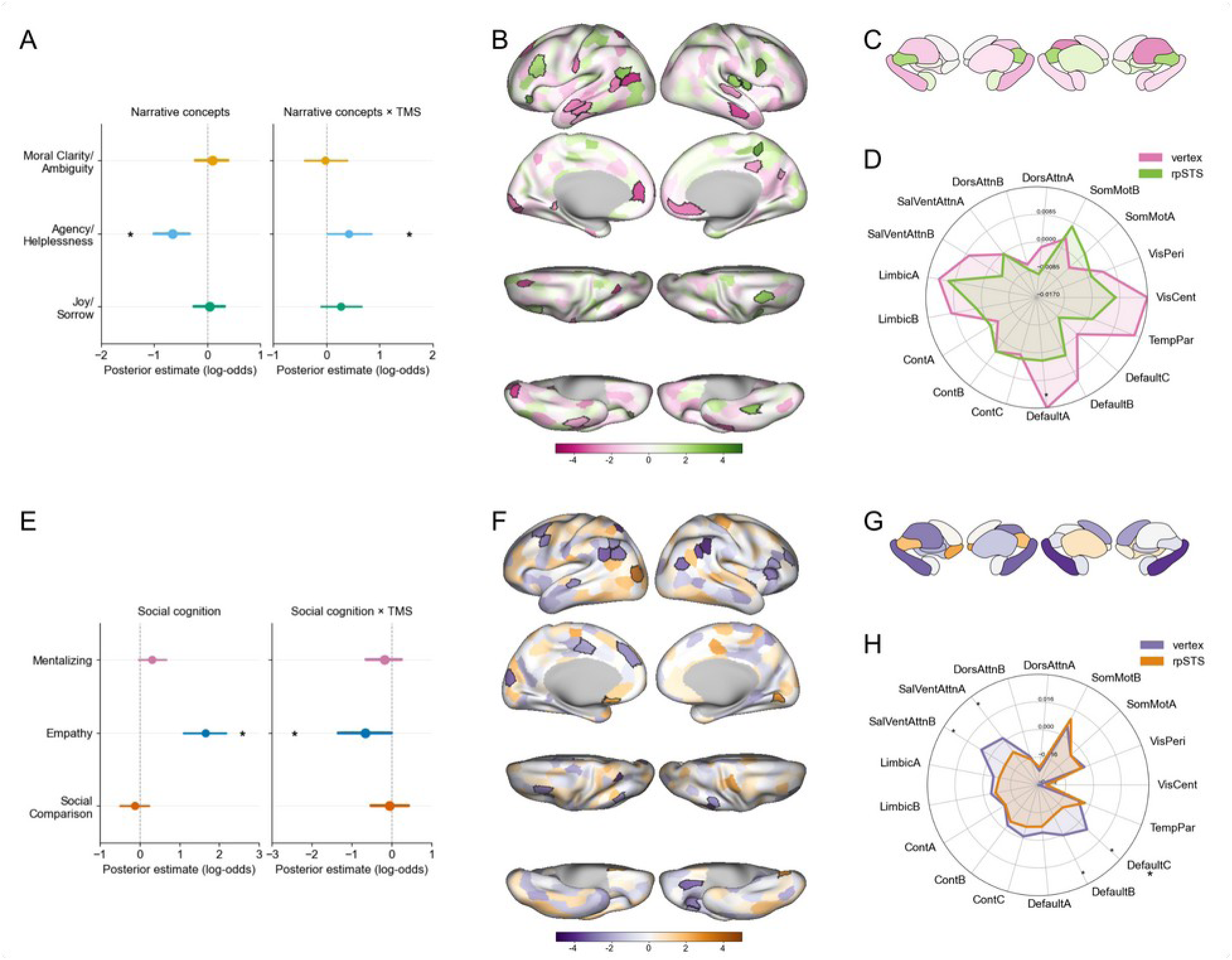
Reduced encoding of social concepts central to the narrative following rTMS to the rpSTS. **A.** Plots represent the posterior mean and 95% HDI estimates of the effects of narrative element concepts and rTMS on the probability of scene recall. The main effect of narrative element concepts is shown on the left, with the interaction between narrative element concepts and stimulation group on the right. Asterisks indicate that the 95% HDI did not contain zero. **B & C.** Group differences in the difference in ISPS between high and low helplessness scenes. The t-values of the comparison between the rpSTS and vertex groups in ISPS from high – low helplessness scenes are shown, with lower ISPS difference in the rpSTS relative to vertex shown in pinks, and higher ISPS difference in the rpSTS relative to vertex group shown in yellow/greens. Parcels are overlaid on the fsaverage6 cortical surface **(B)** or the Melbourne Subcortex Atlas (**C**). Parcels outlined in black or overlaid with an asterisk are p<0.05 (FDR corr). **D.** Data from the 400 cortical parcels collapsed into 17 networks, showing mean difference in ISPS between high and low helplessness scenes for the rpSTS group (yellow/green) and the vertex group (pink). Asterisks indicate significant differences between the groups, p<0.05 (FDR corr). **E.** Plots represent the posterior mean and 95% HDI estimates of the effects of social cognition concepts and rTMS on the probability of scene recall. The main effect of social cognition concepts is shown on the left, with the interaction between narrative elements and stimulation group on the right. Asterisks indicate that the 95% HDI did not contain zero. **F & G.** Group differences in the difference in ISPS between high and low empathy scenes. The t-values of the comparison between the rpSTS and vertex groups in ISPS from high – low empathy scenes are shown, with lower ISPS difference in the rpSTS relative to vertex shown in purples, and higher ISPS difference in the rpSTS relative to vertex group shown in oranges. Parcels are overlaid on the fsaverage6 cortical surface **(F)** or the Melbourne Subcortex Atlas (**G**). Parcels outlined in black or overlaid with an asterisk are p<0.05 (FDR corr). **H.** Data from the 400 cortical parcels collapsed into 17 networks, showing mean difference in ISPS between high and low empathy scenes for the rpSTS group (purple) and the vertex group (orange). Asterisks indicate significant differences between the groups, p<0.05 (FDR corr).

To examine the contribution of the agency/helplessness concept to shared neural representations, we identified the scenes with the highest and lowest helplessness concept weights, as scenes loaded predominantly on the negative pole of this concept. We then calculated the difference in ISPS from high and low helplessness scenes for each group and compared across groups. We observed a greater difference in ISPS to high versus low helplessness scenes in the vertex group compared to the rpSTS group in bilateral medial prefrontal, anterior temporal, and superior temporal cortices, as well as right medial parietal cortex (p<0.05, FDR corr; Figure 5B,C). Greater ISPS to high versus low helplessness scenes following rpSTS stimulation compared to vertex was observed in bilateral dorsolateral prefrontal and right posterior insula cortices. At the network level, greater ISPS for high versus low helplessness scenes in the vertex group compared to the rpSTS group was observed in Default-A subnetwork (p<0.05, FDR corr; Figure 5D).

The movie narrative concepts analyzed above provide details about the semantic contents of the central movie narrative, but these concepts are only indirectly associated with psychological constructs. If one role of shared representations is to assist in making social inferences and understanding – in this case, regarding the characters in the movie – then we might expect that participants encode naturalistic information in a manner that aligns with social-cognitive constructs. To test this idea, we generated a new set of concept embeddings based on concepts drawn from the social cognition literature. In particular, we focused on the following concepts: mentalizing/non-mentalizing, self-other overlap/distinctiveness, social comparison/self comparison, empathy/self absorption, and shared reality/private experience. We examined if these concepts were connected to scene centrality and scene recall. As with the previous analysis, we generated phrases associated with these concepts, created concept embeddings, and then derived concept weights for each scene by examining the similarity of scene embeddings to the concept embeddings. Of the five concepts, only mentalizing/non-mentalizing, posterior mean *b* = 0.77, 95% HDI = [0.44, 1.11], social comparison/self comparison posterior mean *b*=-0.78, 95% HDI = [-1.12, -0.48], and empathy/self absorption, posterior mean *b*=0.64, 95% HDI = [0.22, 1.03], were associated with scene centrality. When we examined if the concept weights were associated with scene recall, we found that only empathy/self absorption was associated with recall, posterior mean *b*=1.66, 95% HDI = [1.01, 2.18]. Overall scenes that were more positive on the empathy weight were recalled better. The empathy content of a scene also showed a stimulation by concept weight interaction for the probability of scene recall, posterior mean *b*=-0.65, 95% HDI = [-1.33, -0.01] (Figure 5E).

As was the case with the agency/helplessness concept, these results indicate that the rpSTS group encoded information in scenes related to empathy less during movie encoding than did the vertex group. We compared ISPS from high versus low empathy scenes, as all scenes loaded on the positive pole of this concept. We observed greater difference in ISPS to high versus low empathy scenes in the vertex group compared to the rpSTS group in bilateral TPJ, right inferior frontal and orbitofrontal cortices, right hippocampus, and left dorsomedial and dorsolateral prefrontal cortices (p<0.05, FDR corr; Figure 5F,G). Examining these data at the network level indicated greater ISPS to high empathy versus low empathy scenes in the vertex group compared to the rpSTS group in Default-B and Default-C subnetworks, as well as Salience/Ventral Attention-A and Salience Ventral Attention-B subnetworks (Figure 5H).

## Discussion

Sharing representations of the world with others can help us develop and strengthen our social connections. The similarity of neural representations in default mode regions, especially during naturalistic movie viewing, is thought to reflect in part the neural basis of these shared representations [8,9]. We used continuous theta-burst rTMS to disrupt neural activity in the rpSTS, and showed that, relative to stimulation of a vertex location, there was lower neural similarity during movie watching in limbic and DMN regions during scenes central to the movie narrative versus noncentral scenes. Using NLP to analyze the main semantic concepts of the movie narrative, we showed that disruption of rpSTS was also associated with decreased memory for central scenes and social details from the movie, relative to vertex stimulation. Our findings provide causal evidence that the rpSTS supports the encoding of social information from the environment into a shared representation in the DMN.

A key finding from our study was that the encoding of scenes higher in centrality (i.e., scenes more closely related to the broader movie narrative) was reduced in participants who received rTMS to the rpSTS when compared to participants in the vertex group. Previous research has shown that semantic centrality is associated with scene memorability, and has suggested that centrality contributes to shared representations in DMN regions [28]. We also observed greater neural pattern similarity between participants in DMN and limbic network regions, including temporoparietal junction, rpSTS, medial prefrontal and orbitofrontal cortices for high centrality vs low centrality scenes in the vertex group, compared to the rpSTS group. In particular, all three subnetworks of the DMN showed greater ISPS for high vs low centrality scenes in the vertex group compared to the rpSTS group. Our results suggest that the rpSTS plays a key role in encoding semantically central elements of a narrative into brain-wide shared representations of central narrative themes.

Centrality itself, however, does not provide details about the content of the movie narrative. To characterize the semantic contents of each scene, we developed a novel NLP-based approach to identify concepts associated with the central narrative. We projected scene transcript embeddings onto concept embeddings drawn from film/narrative theory and social-cognitive psychology literatures. We found that the concepts of moral clarity/moral ambiguity, agency/helplessness, hope/despair (from film/narrative theory), and mentalizing/non-mentalizing, social comparison/self comparison, and empathy/self absorption (from social cognitive psychology), were associated with scene centrality. Only agency/helplessness and empathy/self absorption were linked to scene recall, however, and for both of these concepts the contribution to recall was lower in the rpSTS group than it was in the vertex group. We also found that high vs low helplessness scenes were associated with greater ISPS in the DMN-A subnetwork for the vertex group than for the rpSTS group. High vs low empathy scenes were associated with greater ISPS in the DMN-B and DMN-C subnetworks for the vertex group compared to the rpSTS group. The concept embeddings used in this study do not exhaust the space of possible narrative concepts, and the concepts correlated with scene centrality here might be specific to this movie. However, these analyses provide valuable insight into the narrative content whose encoding is disrupted by rTMS to the rpSTS, and may provide an approach to further understand narrative processing in naturalistic stimuli.

Disruption of the encoding of scenes high in helplessness and empathy via rTMS to the rpSTS is consistent with the role of this region in empathic cognition [32,33]. The rpSTS has been suggested to be a key region in a social perception pathway that runs parallel to the object (“what”) pathway and the spatial (“where”) pathway [24]. This region receives extensive inputs from motion-selective and human form-selective regions upstream in the visual pathway, and prior TMS studies have shown that disruption of the rpSTS impairs the perception of dynamic social information [25,34]. But there is also strong evidence that this region is not just a high-level visual region, and instead interacts with downstream regions for the conceptual processing of social information, including regions in the DMN [35–37]. Our findings highlight the important role that rpSTS plays in shared representations within these regions during naturalistic viewing. Our results suggest that the rpSTS acts as a key hub in the integration of social perceptual processing with higher-order social cognition.

In contrast to the effects of rpSTS disruption on neural similarity in regions in the DMN and limbic networks, we observed greater synchrony and neural pattern similarity in early visual regions and frontal cortex in the rpSTS group compared to the vertex group. In particular, we found that ISPS in left dorsolateral prefrontal cortex to high vs low centrality scenes was greater in the rpSTS group than in the vertex group. The left dorsolateral prefrontal cortex is a key region for episodic memory encoding [38,39]. These neural findings are consistent with the behavioral findings that the rpSTS group showed a difference in recognition memory for non-social vs social details that the vertex group did not, and overall did not show lower scene recall than the vertex group. We suggest that rather than producing a general encoding deficit, disruption of the rpSTS resulted in a shift in encoding away from central, social details and towards more peripheral aspects of the movie.

Together, our findings establish a role for the rpSTS in encoding central, socially relevant narrative details during naturalistic viewing into a shared neural representation that is broadcast throughout the DMN and limbic network. These results support the proposal that the rpSTS is an important interface between the visual stream processing social stimuli and higher order networks that underlie complex social cognition. More broadly, our findings show that shared representations in networks such as the DMN rely on the encoding of inputs from the external environment, and suggest that when such inputs are disrupted neural patterns within DMN become more idiosyncratic.

## Methods

### Participants

In this study, 44 participants (ages 18-39 years, mean age=22.2, S.D.=4.4, 33F) completed an initial screening session and two sessions involving fMRI and TMS. Participants were required to be right-handed, have normal or corrected-to-normal vision, be native/fluent English speakers, and have no history of neurological or psychiatric disorders that would exclude them from receiving TMS. The initial screening session took 30min, and the two fMRI and TMS sessions each took ∼1h. Participants were paid $50 upon completion of all three sessions. Participants provided written informed consent, and all procedures were approved by the George Mason University Human Subjects Review Board.

### Procedure

During initial screening, participants first provided written consent and completed a demographics survey. Participants were then interviewed by a researcher to determine if they met any inclusion/exclusion criteria for MRI and for administration of TMS. Participants who met inclusion criteria for MRI and TMS then attended two sessions.

In the first MRI and TMS session, resting motor thresholds (rMTs) were obtained from single pulses to the hand area of left motor cortex using a MagPro x100 stimulator and C-B60 coil (MagVenture). rMT was defined as the lowest stimulator output needed to evoke 5 visible finger movements from 10 pulses, with a starting stimulator output of 55%. Following rMTs, participants were placed in the MRI scanner and BOLD EPI data were collected while participants viewed a localizer task. The localizer task consisted of 3sec videos of people (mostly smiling and interacting in groups), places (natural scenes of pleasant outdoor locations), objects (high value objects such as money, jewels, luxury cars), food (appetizing foods such as chocolate and french fries), and scrambled versions of videos from all categories. Videos from each category were presented in 16sec blocks (5 videos, 200msec ISI) in pseudorandomized order, with 3 blocks of each category per run and 2 runs per participant. Each run had 2 16sec rest periods randomly interspersed among the video blocks, as well as 16sec rest periods at the start and end of the run. Participants were asked to identify when videos within a block repeated. High resolution T1w anatomical images were also acquired. BOLD fMRI data were preprocessed and analyzed (see Imaging Preprocessing and Imaging Analysis) and the contrast of people > all other categories at z>4.0 (two-tailed p<0.000064, uncorrected) used to identify the rpSTS in each participant’s native T1w space. All but one participant showed a peak response in the rpSTS at z>4.0, with the one participant showing a peak response in the rpSTS at z>3.0.

The second MRI and TMS session occurred between 2 and 7 days after the first session. In this session, participants were randomly allocated to one of two groups, and administered continuous theta burst stimulation (3 pulse burst of 50Hz at 5hz, 600 pulses, 70% of rMT) to either the rpSTS or a vertex location. Coil location for stimulation was guided using Brainsight (Rogue Research). Following the end of stimulation, participants were moved into the MRI scanner, where they then watched the 20min movie “The Neighbors Window” [40]. Movie presentation and scanning started within 3min of the completion of cTBS in all participants. Once the movie was complete, participants were removed from the scanner and performed a task where they freely recalled the events of the movie and then answered a series of recognition questions. Free recall responses were recorded using Audacity.

#### Movie annotations and ratings

Three annotators separately viewed the movie in three-second sections and provided written descriptions of the sections. In a separate session, the three annotators rated each three second segment for arousal, valence, and communicative interactions. All three annotators then jointly segmented the movie into distinct scenes with scene start and end times using consensus coding. Transition scenes consisting of a blank screen were dropped from further scene-level analysis.

#### Movie features extraction

Features describing the visual, auditory and speech content of the film were extracted at high temporal resolution and reduced to the TR sampling rate. All feature extraction was performed in Python 3.12 on the same video file that was presented to participants.

#### Faces and bodies

Face and body features were extracted using the MediaPipe Tasks API (version 1.0.1; Google). Every third movie frame was analyzed. Faces were detected with the BlazeFace short-range model, from which we computed the number of detected faces per frame and the summed face bounding-box area as a proportion of total frame area. Body pose was estimated with the Pose Landmarker (full model) configured to detect up to four people per frame, returning 33 landmarks per person in normalized image coordinates. The number of detected people was recorded per frame. Body velocity was computed as the mean inter-frame displacement of landmarks, divided by the inter-frame interval and averaged across detected people. People were matched between consecutive sampled frames by nearest centroid (maximum displacement 0.25 normalized units) to limit identity-swap artifacts.

#### Motion energy

Low-level visual motion energy was computed with pymoten version 0.1.3 [41]. Frames were converted to grayscale, resized to 96 × 96 pixels, and resampled to 10 Hz by nearest-timestamp selection. The resulting sequence was projected through the default spatiotemporal Gabor filter pyramid (1380 filters spanning five spatial frequencies of 0–16 cycles per image, three temporal frequencies of 0–4 Hz, and eight directions of motion at 45° intervals; filter temporal width 6 frames). Filter outputs were log-transformed and averaged across channels to yield a single motion-energy timecourse.

#### Auditory amplitude

Audio was extracted with ffmpeg as mono 22.05 kHz PCM and analyzed with librosa version 1.0.0 [42]. A short-time Fourier transform (1024-point window) yielded a broadband root-mean-square amplitude envelope.

#### Speech

Speech was detected using with faster-whisper version 1.2.1 using (small.en model) with word-level timestamps and voice-activity filtering enabled. A speech regressor was formed as the proportion of each 1.5s time bin overlapped by transcribed utterance intervals, so that pauses within an utterance count as speech.

#### Annotation embeddings

Each set of annotations from the three-second sections from each annotator was converted to 768-dimensional sentence embeddings using SBERT (all-mpnet-base-v2). Embeddings from each scene were L2-normalized then averaged together. The mean cosine similarity of embeddings across the three annotators = 0.80 (average s.d. = 0.16). scene-level embeddings were then averaged across the three annotators.

### Behavior

Accuracy and reaction time for social and nonsocial items were collected from the recognition memory test. Responses associated with reaction times less than 50ms or more than 2500ms were excluded from further analysis. Participant verbal recall was transcribed to text using OpenAI Whisper implemented in a Google Colab notebook and then manually checked. Recall transcripts were then segmented into phrases based on pauses and changes in topic.

#### Recall scoring

Two of the three annotators rated each recall segment as reflecting recall of a scene, using a method based on [43]. A scene was scored as recalled if a recall segment described any part of the scene, and a segment could be assigned to more than one scene.

### Statistical analyses of behavior were conducted in R v4.6.0 using brms

v2.23.0/CmdStan v2.39.0. All analyses used 4 chains, No-U-Turns sampler (NUTS), 2000 iterations per chain with 1000 iteration burn in, seed=42, adapt_delta = 0.95 or 0.99. Inferences were based on the 95% highest density interval (HDI). Dependent variables were modeled with appropriate distributions: beta regression for proportions bounded on (0, 1), Gaussian regression for continuous ratio-scaled variables (e.g., log(RT)), and Bernoulli regression with a logit link for binary outcomes (e.g., recall accuracy). Bivariate correlations between concept weights and centrality were examined using robust (student’s t) regression to reduce the impact of outliers. Unless otherwise noted, weakly informative priors were used: Normal (0,5) for beta and Gaussian or Normal(0, 2.5) for Bernoulli models for the intercept and fixed-effect coefficients, and Exponential(1) for the standard deviation of participant-level random effects. Model comparison used approximate leave-one-out cross-validation (LOO-CV) [44] and expected log predictive density difference (elpd_diff).

### Scene Networks and Concept Embedding

Pairwise similarity between scene embeddings was computed as the dot product between each pair of normalized embedding vectors, yielding an n × n similarity matrix (where n is the number of scenes). Degree centrality for each scene was calculated as the sum of its similarities to all other scenes and normalized by dividing by n − 1, yielding a centrality value bounded between 0 and 1 that reflects the average similarity of a given scene to all other scenes in the set.

To examine the semantic content of the narrative, we projected each scene embedding onto a set of concept axes (see below). Following [30], concept axes were constructed by encoding positive and negative pole descriptor phrases using the same sentence embedding model (SBERT all-mpnet-base-v2) used to embed scene transcripts. The phrases associated with positive and negative poles of each concept were generated using Claude Sonnet 4.6 (Anthropic) (Supplementary Methods). The sampling parameters are not user-configurable in the web interface, so phrase generation is not deterministically reproducible; the full set of generated phrases is provided below and all reported analyses use the fixed set. The concepts (and associated phrases) are not exhaustive and were chosen as they reflected some of the key narrative components of the movie.

Movie Narrative Concept Axes: Concept axes were constructed for seven dimensions drawn from narrative film theory [45–47]:

#### Danger/Safety

The degree to which a scene reflects threat and danger versus security and calm. Positive pole: danger threat imminent peril dread fear menace stalking; Negative pole: safety calm security comfort shelter peaceful tranquil protected.

#### Chaos/Order

The extent to which a scene reflects disorder vs order. Positive pole: disorder disruption breakdown anarchy disintegration collapse confusion unravel; Negative pole: order structure control stability equilibrium routine predictable organized. <u>Joy/Grief: T</u>he emotional valence of a scene. Positive pole: happiness celebration laughter delight elation euphoria cheerful festive; Negative pole: grief sorrow mourning loss devastation weeping heartbreak despair.

#### Transformation/Stasis

Whether a scene is oriented toward change and growth versus repetition and fixedness. Positive pole: change growth evolution turning point breakthrough metamorphosis becoming new; Negative pole: stasis repetition static fixed unchanged trapped loop regression same.

#### Moral Clarity/Moral Ambiguity

Whether a scene presents a clear moral stance versus moral complexity. Positive pole: righteousness clear good evil defined justice heroic noble virtuous; Negative pole: ambiguous morally complex uncertain ethical doubt gray complicit corrupt.

#### Agency/Helplessness

The degree to which characters exercise goal-directed control versus passiveness. Positive pole: action decision control pursuit determination drive willpower initiative; Negative pole: helplessness passivity victimhood powerless trapped paralyzed submission surrender.

#### Hope/Despair

Future-oriented affective orientation in a scene. Positive pole: optimism possibility future belief aspiration faith potential bright horizon; Negative pole: despair hopeless futile doom defeat resignation bleakness giving up.

### Social Cognition Concept Axes

To examine the social cognitive dimensions of scene content, we constructed a second set of concept axes drawn from the social cognition literature. These axes captured dimensions of social cognition and inference relevant to the movie’s central narrative.

#### Mentalizing/Non-mentalizing

The degree to which a scene is oriented toward inferring the inner beliefs, intentions, and feelings of others, versus attending only to observable surface behavior and external appearance. [48,49]. Positive pole descriptors: understanding minds beliefs intentions feelings perspective taking inner life. Negative pole descriptors: surface behavior appearance observable action external ignoring inner states.

#### Self-other overlap/distinctiveness

The degree to which another person is incorporated into one’s own self-concept versus perceiving the other as a separate and autonomous individual [50,51]. Positive pole descriptors: identification merging with other seeing yourself in them projection mirroring. Negative pole descriptors: other as separate distinct different autonomous independent bounded self.

#### Social comparison/self comparison

Whether scene content is oriented toward evaluating oneself relative to others versus a self-referential orientation that is not concerned with status relative to others [52,53]. Positive pole descriptors: comparing upward comparison ranking status measuring against others better worse. Negative pole descriptors: self-sufficient not comparing autonomous self-referential unconcerned with others.

#### Shared reality/Private Experience

The degree to which inner experience is communicated and mutually understood between characters versus inside and invisible to others [1]. Positive pole descriptors: communicated mutual understanding shared expressed visible known to others. Negative pole descriptors: private hidden locked inside invisible interior unspoken unknowable alone.

#### Empathy/Self-absorption

The degree to which scene content is oriented toward compassion, feeling the same as others, and other-directed care, versus self-absorption and inward preoccupation making one oblivious to the experience of others [54]. Positive pole descriptors: compassion understanding fellow feeling grief for another perspective taking care. Negative pole descriptors: self-absorption self-pity inward preoccupation oblivious to others focused on own problems.

We computed the mean embedding for each pole, and then subtracted the negative pole mean embedding from the positive pole mean embedding to yield a unit-normalized direction vector in 768-dimensional embedding space. To project each scene embedding onto a concept axis, we calculated the cosine similarity between the scene embedding and the concept direction to derive concept weights for each scene, with weights that range from −1 (fully aligned with the negative pole) to +1 (fully aligned with the positive pole).

#### Imaging acquisition

Structural and functional MRI data were collected using a 3T Siemens Prisma MRI scanner and a 32 channel phased array head coil located at George Mason University. BOLD fMRI data were collected using a multiband, multiecho gradient echo T2*-weighted echoplanar imaging (EPI) sequence (TR=1.5s, TE=14/33/52ms, GRAPPA acceleration factor=2, multiband acceleration factor=3, flip angle=68°, voxel size=2.7mm^3^, axial slices). A dual-echo gradient recalled echo (GRE) fieldmap (TR=0.73s, TE=4.9/7.3ms, flip angle=50°, voxel size=2.7mm^3^, axial slices) was also collected to correct for susceptibility-based distortion. A high resolution T1w MPRAGE anatomical scan (TR=2.4s, TE=2.4ms, GRAPPA acceleration factor=2, flip angle=8°, voxel size=0.8mm^3^, sagittal slices) was acquired for coregistration and cortical surface-based analyses, and a larger FOV T1w MPRAGE anatomical scan (TR=1.2s, TE=2.6ms, flip angle=12°, voxel size=1.0mm^3^, axial slices) was acquired for MRI guided neuronavigation of the TMS coil.

#### Imaging preprocessing

Results included in this manuscript come from preprocessing performed using fMRIPrep 23.1.4 [55,56], which is based on Nipype 1.8.6 [57,58], and tedana 0.0.12 [59]. T1w anatomical scans were corrected for intensity non-uniformity. Brain surfaces were reconstructed from the high resolution sagittal T1w image using FreeSurfer. Volume-based spatial normalization to the MNI152NLin2009cAsym standard space was performed through nonlinear registration with antsRegistration (ANTs). Each of the BOLD runs was corrected for susceptibility distortion using fieldmaps and motion corrected. BOLD data were coregistered to a T1w anatomical image. TE-dependence analysis was performed on fMRIPrep preprocessed input data using the tedana workflow. Independent components analysis with component selection was performed to identify BOLD (TE-dependent), non-BOLD (TE-independent), and uncertain (low-variance) components.

#### fMRI analysis

Statistical analysis of the localizer data was performed using FSL’s FEAT [60]. The optimally combined and denoised (optcomDenoised) TE-dependent data in axial T1w space from tedana preprocessing for each of the two runs were smoothed with a 6mm FWHM gaussian kernel. Data were temporally filtered at 1/128s and prewhitened using FILM, and a GLM that included regressors for each of the five conditions, as well as covariates for the first four TRs and for TRs in which the framewise displacement from the bandstop filtered motion parameters exceeded 0.5mm [61] was estimated. Contrasts comparing the people blocks vs the average of the other four conditions were averaged across the two runs in a fixed effects analysis.

The voxel timeseries of optimally combined and denoised (optcomDenoised) TE-dependent movie data in sagittal T1w space from tedana preprocessing were converted to zscores. TRs in which the framewise displacement from the bandstop filtered motion parameters exceeded 0.5mm were censored and missing data interpolated using Lomb-Scargle interpolation [62]. Data from three participants (two from vertex group, one from rpSTS group) were excluded from further analyses due to excessive motion. Data were projected to the fsaverage6 cortical surface (40,962 vertices per hemisphere) [63] using FreeSurfer v6.0.0 and smoothed on the surface with a 2mm FWHM gaussian kernel. To facilitate subcortical parcellation, data were also warped to the MNI152NLin2009cAsym atlas using ANTs and smoothed with a 2mm FWHM gaussian kernel. Right and left hemisphere surface data were parcellated using the 400 parcel Schaefer atlas [64]. Subcortical data were parcellated using the 16 parcel Melbourne Subcortex S1 Atlas [65].

To fit the encoding model, we first extracted movie timeseries from rpSTS and EVC ROIs from participants from both stimulation groups. The location of the rpSTS in T1w space of each participant was projected onto their cortical surface and an 8mm radius circular ROI label was made. The EVC ROI was constructed from the FreeSurfer V1_exvivo.thresh label. Preprocessed movie timeseries were extracted from these labels. The seven model features (number of faces, number of bodies, body motion velocity, speech segments, observer ratings of communicative interactions, motion energy, and rms envelope) were resampled to the TR and convolved with a double-gamma hemodynamic response function. The variance inflation factors of the features of the model were all below 2.0. The model was fit to the rpSTS and EVC ROI timeseries using cross-validated (CV) ridge regression. Timeseries were split into six segments of equal length; five of the six segments were used to tune the ridge regularization parameter using five-fold CV, and then used to train the model. Model prediction performance was then tested on the sixth held-out segment. Timepoints in the held-out segment within 10 TRs from a training set segment were dropped to reduce leakage between training and test data due to long-range autocorrelation. This process was repeated six times.

Model performance was evaluated via the Pearson correlation between the predicted and observed timeseries in the held-out segments, averaged across repetitions after Fisher z-transformation. Unique variance explained for each group of features (social, auditory, visual) was calculated following [66] as:

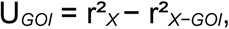

where X is the full feature set, GOI is the feature group of interest, and r² the squared held-out prediction correlation. Each reduced model was refit through the same nested procedure with its own regularization parameter selected by the inner cross-validation. We made one addition to this measure. Because r² is always positive, a model predicting in the wrong direction will appear as explaining variance no different from one predicting in the correct direction; where prediction is near chance this can inflate apparently unique contributions. We therefore multiplied each term by the direction of its correlation, sign(r)·r², which is identical to the original measure whenever prediction is above chance and centers cases of null correlation to zero. Fisher’s z-transformed *r* values and signed U*_GOI_* values from the different models, ROIs, and stimulation groups were then compared using linear mixed effects models in brms.

To calculate intersubject correlations, for each participant in each group we calculated the average fMRI time series within each of the 416 parcels, and then calculated the Pearson correlation using scipy.stats [67] between time series from a given parcel and the corresponding parcels from every other participant in their group. To calculate the intersubject pattern similarity, we first segmented the fMRI time series into the previously identified scenes. We then calculated the average spatial pattern for each vertex or voxel for each scene, and then for each participant calculated the Pearson correlation between spatial patterns for a given parcel and corresponding parcels from every other participant in their group. The ISC and ISPS values were then Fisher’s z-transformed. For the analysis of overall ISPS, we averaged ISPS across scenes. For the comparison of high versus low centrality scenes, we subtracted the mean ISPS from the bottom 40% centrality scenes from the mean ISPS from the top 40% centrality scenes. For the comparison of high versus low helplessness scenes, we subtracted the mean ISPS from the bottom 40% helplessness (i.e., top 40% agency) scenes from the mean ISPS from the top 40% helplessness (i.e., bottom 40% agency) scenes. For the comparison of high versus low empathy scenes, we subtracted the mean ISPS from the bottom 40% empathy scenes from the mean ISPS from the top 40% empathy scenes. For network-based analyses, ISC and ISPS values were averaged within each of the 17 networks of the Yeo parcellation [68]. Group differences in ISC and ISPS were calculated using t-tests for independent samples and p-values calculated from permutation tests with 5000 permutations, along with false discovery rate correction, using scipy.stats.

## Code availability

Data analyses and code associated with this study can be found at https://github.com/CompSocNeuroGroup/rTMS_narrative_concepts

## Data availability

Data from this study will be made available upon publication at OpenNeuro.

## Acknowledgments

The authors thank Sima El-Bash and Michalina Pilch for assistance with data collection, and Martin Wiener for assistance with TMS and for thoughtful comments about the manuscript.

## Supplementary Methods

The concept weights for each scene were derived by projecting the scene embeddings onto bipolar concept axes. Each axis was defined by two sets of descriptor phrases, one per pole, generated with an LLM. Details on the model and prompts are provided below.

Model and access conditions. Claude Sonnet 4.6 (Anthropic) was used via the claude.ai web application in May 2026. Each prompt was delivered in a new conversation with no prior history. Extended thinking was not enabled and websearch was not recorded. Sampling temperature was not fixed as the phrases were generated through the website rather than the API, and so deterministic reproduction of the generated phrases is not possible. The generated phrase sets are provided in full, and all reported analyses were run on the fixed phrase sets.

Prompt 1: Film and narrative theory concepts

“Do not use any previous session history in this session.

I am constructing semantic concept axes for a study of narrative film

comprehension. The film is “The Neighbors Window” by Marshall Curry, which is based on “The Living Room”, a true story recounted by Diane Weipert on the podcast Love and Radio.

To study each scene in the movie, I need to construct bipolar concept axes by defining positive and negative poles using short descriptor phrases. Each pole should be 6–8 single words or very short phrases separated by spaces, not prose.

The concepts are:

Danger/Safety: The degree to which a scene reflects threat and danger versus security and calm.

Chaos/Order: The extent to which a scene reflects order versus disorder.

Joy/Grief: The emotional valence of a scene.

Transformation/Stasis: Whether a scene is oriented toward change and growth versus repetition and fixedness.

Moral Clarity/Ambiguity: Whether a scene presents a clear moral stance versus moral ambiguity.

Agency/Helplessness: The degree to which characters exercise goal-directed control versus passiveness.

Hope/Despair: Future-oriented emotional orientation in a scene.

Return the concepts and phrases in a python dictionary format saved as a .txt file.”

Prompt 2: Social cognition concepts

“Do not use any previous session history in this session.

I am constructing semantic concept axes for a study of social cognition concepts in scenes from a film. The film is “The Neighbors Window” by Marshall Curry, which is based on “The Living Room“, a true story recounted by Diane Weipert on the podcast Love and Radio.

To study each scene in the movie, I need to construct bipolar concept axes by defining positive and negative poles using short descriptor phrases. Each pole should be 6–8 single words or very short phrases separated by spaces, not prose. These are technical constructs from the social cognition literature, not everyday terms. Use the definitions given below rather than lay meanings.

Mentalizing/non-mentalizing: The degree to which a scene is oriented toward inferring the inner beliefs, intentions, and feelings of others, versus attending only to observable surface behavior and external appearance.

Self-other overlap/distinctiveness: The degree to which another person is incorporated into one’s own self-concept versus perceiving the other as a separate and autonomous individual.

Social comparison/self comparison: Whether scene content is oriented toward evaluating oneself relative to others versus a self-referential orientation that is not concerned with status relative to others.

Shared reality/private experience: The degree to which inner experience is communicated and mutually understood between characters versus inside and invisible to others.

Empathy/self-absorption: The degree to which scene content is oriented toward compassion, feeling the same as others, and other-directed care, versus self-absorption and inward preoccupation making one oblivious to the experience of others.

Return the concepts and phrases in a python dictionary format saved as a .txt file.”

## References

1. Echterhoff, G., Higgins, E. T. & Levine, J. M. Shared reality: experiencing commonality with others’ inner states about the world. Perspect. Psychol. Sci. 4, 496–521 (2009).

2. Wolf, W. & Tomasello, M. A shared intentionality account of uniquely human social bonding. Perspect. Psychol. Sci. 20, 264–275 (2025).

3. Rossignac-Milon, M. & Higgins, E. T. Epistemic companions: shared reality development in close relationships. Curr. Opin. Psychol. 23, 66–71 (2018).

4. Nguyen, M., Vanderwal, T. & Hasson, U. Shared understanding of narratives is correlated with shared neural responses. Neuroimage 184, 161–170 (2019).

5. Baek, E. C. & Parkinson, C. Shared understanding and social connection: integrating approaches from social psychology, social network analysis, and neuroscience. Soc. Personal. Psychol. Compass 16, e12710 (2022).

6. Parkinson, C., Kleinbaum, A. M. & Wheatley, T. Similar neural responses predict friendship. Nat. Commun. 9, 332 (2018).

7. Shen, Y. L. et al. Neural similarity predicts whether strangers become friends. *Nat*. Hum. Behav. 9, 2285–2298 (2025).

8. Yeshurun, Y., Nguyen, M. & Hasson, U. The default mode network: where the idiosyncratic self meets the shared social world. Nat. Rev. Neurosci. 22, 181–192 (2021).

9. Ohad, T., Zvi, Y. & Yeshurun, Y. Default mode network synchrony reflects shared understanding. Curr. Opin. Behav. Sci. 64, 101540 (2025).

10. Finn, E. S. et al. Idiosynchrony: from shared responses to individual differences during naturalistic neuroimaging. Neuroimage 215, 116828 (2020).

11. Lahnakoski, J. M. et al. Naturalistic fMRI mapping reveals superior temporal sulcus as the hub for the distributed brain network for social perception. Front. Hum. Neurosci. 6, 233 (2012).

12. Simony, E. et al. Dynamic reconfiguration of the default mode network during narrative comprehension. Nat. Commun. 7, 12141 (2016).

13. Allison, T., Puce, A. & McCarthy, G. Social perception from visual cues: role of the STS region. Trends Cogn. Sci. 4, 267–278 (2000).

14. Seltzer, B. & Pandya, D. N. Afferent cortical connections and architectonics of the superior temporal sulcus and surrounding cortex in the rhesus monkey. Brain Res. 149, 1–24 (1978).

15. Seltzer, B. & Pandya, D. N. Intrinsic connections and architectonics of the superior temporal sulcus in the rhesus monkey. J. Comp. Neurol. 290, 451–471 (1989).

16. Seltzer, B. & Pandya, D. N. Frontal lobe connections of the superior temporal sulcus in the rhesus monkey. J. Comp. Neurol. 281, 97–113 (1989).

17. Seltzer, B. & Pandya, D. N. Post-rolandic cortical projections of the superior temporal sulcus in the rhesus monkey. J. Comp. Neurol. 312, 625–640 (1991).

18. Baizer, J. S., Ungerleider, L. G. & Desimone, R. Organization of visual inputs to the inferior temporal and posterior parietal cortex in macaques. J. Neurosci. 11, 168–190 (1991).

19. Boussaoud, D., Ungerleider, L. G. & Desimone, R. Pathways for motion analysis: cortical connections of the medial superior temporal and fundus of the superior temporal visual areas in the macaque. J. Comp. Neurol. 296, 462–495 (1990).

20. Felleman, D. J. & Van Essen, D. C. Distributed hierarchical processing in the primate cerebral cortex. Cereb. Cortex 1, 1–47 (1991).

21. Turner, B. H., Mishkin, M. & Knapp, M. Organization of the amygdalopetal projections from modality-specific cortical association areas in the monkey. J. Comp. Neurol. 191, 515–543 (1980).

22. Edmonds, D. et al. The human social cognitive network contains multiple regions within the amygdala. Sci. Adv. 10, eadp0453 (2024).

23. Saygin, A. P. Superior temporal and premotor brain areas necessary for biological motion perception. Brain 130, 2452–2461 (2007).

24. Pitcher, D. & Ungerleider, L. G. Evidence for a third visual pathway specialized for social perception. Trends Cogn. Sci. 25, 100–110 (2021).

25. Basil, R. A., Westwater, M. L., Wiener, M. & Thompson, J. C. A causal role of the right superior temporal sulcus in emotion recognition from biological motion. Open Mind 2, 26–36 (2017).

26. Kensinger, E. A. & Corkin, S. Memory enhancement for emotional words: are emotional words more vividly remembered than neutral words? Mem. Cognit. 31, 1169–1180 (2003).

27. Park, J. S. et al. Emotional arousal enhances narrative memories through functional integration of large-scale brain networks. *Nat*. Hum. Behav. 10, 370–383 (2026).

28. Lee, H. & Chen, J. Predicting memory from the network structure of naturalistic events. Nat. Commun. 13, 4235 (2022).

29. Fenerci, C., Cheng, Z., Addis, D. R., Bellana, B. & Sheldon, S. Studying memory narratives with natural language processing. Trends Cogn. Sci. 29, 516–525 (2025).

30. Kozlowski, A. C., Taddy, M. & Evans, J. A. The geometry of culture: analyzing the meanings of class through word embeddings. Am. Sociol. Rev. 84, 905–949 (2019).

31. Piper, A., So, R. J., & Bamman, D. Narrative theory for computational narrative understanding. In Proceedings of the 2021 conference on empirical methods in natural language processing, 298–311 (2021).

32. Decety, J. & Jackson, P. L. The functional architecture of human empathy. Behav. Cogn. Neurosci. Rev. 3, 71–100 (2004).

33. Zaki, J., Weber, J., Bolger, N. & Ochsner, K. The neural bases of empathic accuracy. Proc. Natl Acad. Sci. USA 106, 11382–11387 (2009).

34. Pitcher, D., Duchaine, B. & Walsh, V. Combined TMS and fMRI reveal dissociable cortical pathways for dynamic and static face perception. Curr. Biol. 24, 2066–2070 (2014).

35. McMahon, E., & Isik, L. Abstract social interaction representations along the lateral pathway. Trends in Cognitive Sciences, 28(5), 392–393 (2024).

36. Miao, Z. et al. Common and distinct neural correlates of social interaction processing and theory of mind in narratives. Nat. Commun. 17, 4830 (2026).

37. Malik, M., Kim, M., Shu, T., Liu, S. & Isik, L. Bottom-up and generative computations uniquely explain neural responses across the social brain. Preprint at bioRxiv 10.64898/2026.02.20.707082 (2026).

38. Cabeza, R. et al. Functional neuroanatomy of recall and recognition: a PET study of episodic memory. J. Cogn. Neurosci. 9, 254–265 (1997).

39. Murray, L. J. & Ranganath, C. The dorsolateral prefrontal cortex contributes to successful relational memory encoding. J. Neurosci. 27, 5515–5522 (2007).

40. Curry, M. The Neighbors’ Window (Marshall Curry Productions, 2019).

41. Nunez-Elizalde, A. O., Deniz, F., Dupré la Tour, T., Visconti di Oleggio Castello, M. & Gallant, J. L. pymoten: scientific python package for computing motion energy features from video. Zenodo 10.5281/zenodo.6349625 (2021).

42. McFee, B. et al. librosa: audio and music signal analysis in Python. In Proc. 14th Python in Science Conference 18–25 (2015).

43. Chen, J. et al. Shared memories reveal shared structure in neural activity across individuals. Nat. Neurosci. 20, 115–125 (2017).

44. Vehtari, A., Gelman, A. & Gabry, J. Practical Bayesian model evaluation using leave-one-out cross-validation and WAIC. Stat. Comput. 27, 1413–1432 (2017).

45. Bordwell, D. Narration in the Fiction Film (Univ. Wisconsin Press, 1985).

46. Todorov, T. & Weinstein, A. Structural analysis of narrative. NOVEL: A Forum on Fiction 3, 70–76 (1969).

47. Shimamura, A. P. (ed.) Psychocinematics: Exploring Cognition at the Movies (Oxford Univ. Press, 2013).

48. Premack, D. & Woodruff, G. Does the chimpanzee have a theory of mind? Behav. Brain Sci. 1, 515–526 (1978).

49. Frith, C. D. & Frith, U. Mechanisms of social cognition. Annu. Rev. Psychol. 63, 287–313 (2012).

50. Aron, A., Aron, E. N. & Smollan, D. Inclusion of Other in the Self Scale and the structure of interpersonal closeness. J. Pers. Soc. Psychol. 63, 596–612 (1992).

51. Decety, J. & Sommerville, J. A. Shared representations between self and other: a social cognitive neuroscience view. Trends Cogn. Sci. 7, 527–533 (2003).

52. Festinger, L. A theory of social comparison processes. Hum. Relat. 7, 117–140 (1954).

53. Fliessbach, K. et al. Social comparison affects reward-related brain activity in the human ventral striatum. Science 318, 1305–1308 (2007).

54. Davis, M. H. Measuring individual differences in empathy: evidence for a multidimensional approach. J. Pers. Soc. Psychol. 44, 113–126 (1983).

55. Esteban, O., et al. fMRIPrep 23.0.2. Zenodo 10.5281/zenodo.852659 (2018).

56. Esteban, O. et al. fMRIPrep: a robust preprocessing pipeline for functional MRI. Nat. Methods 16, 111–116 (2019).

57. Gorgolewski, K. et al. Nipype: a flexible, lightweight and extensible neuroimaging data processing framework in Python. Front. Neuroinform. 5, 13 (2011).

58. Gorgolewski, K. J., et al. Nipype. Zenodo 10.5281/zenodo.596855 (2018).

59. DuPre, E. et al. TE-dependent analysis of multi-echo fMRI with tedana. J. Open Source Softw. 6, 3669 (2021).

60. Woolrich, M. W., Ripley, B. D., Brady, M., & Smith, S. M. Temporal Autocorrelation in Univariate Linear Modeling of FMRI Data. NeuroImage, 14, 1370–1386. (2001).

61. Fair, D. A. et al. Correction of respiratory artifacts in MRI head motion estimates. Neuroimage 208, 116400 (2020).

62. Goldberg, A., Rosario, I., Power, J., Horga, G. & Wengler, K. Strategies for motion- and respiration-robust estimation of fMRI intrinsic neural timescales. Imaging Neurosci. 2, 10.1162/imag_a_00326 (2024).

63. Fischl, B. FreeSurfer. Neuroimage 62, 774–781 (2012).

64. Schaefer, A. et al. Local-global parcellation of the human cerebral cortex from intrinsic functional connectivity MRI. Cereb. Cortex 28, 3095–3114 (2018).

65. Tian, Y., Margulies, D. S., Breakspear, M. & Zalesky, A. Topographic organization of the human subcortex unveiled with functional connectivity gradients. Nat. Neurosci. 23, 1421–1432 (2020).

66. Masson, H. L., & Isik, L. Functional selectivity for social interaction perception in the human superior temporal sulcus during natural viewing. NeuroImage, 245, 118741. (2021).

67. Virtanen, P. et al. SciPy 1.0: fundamental algorithms for scientific computing in Python. Nat. Methods 17, 261–272 (2020).

68. Yeo, B. T. T. et al. The organization of the human cerebral cortex estimated by intrinsic functional connectivity. J. Neurophysiol. 106, 1125–1165 (2011).

